# TKSA-MC: A Web Server for rational mutation through the optimization of protein charge interactions

**DOI:** 10.1101/221556

**Authors:** Vinícius G. Contessoto, Vinícius M. de Oliveira, Bruno R. Fernandes, Gabriel G. Slade, Vitor B. P. Leite

## Abstract

The TKSAMC is a web server which calculates protein charge-charge interactions via the Tanford-Kirkwood Surface Accessibility model with the Monte Carlo method for sampling different protein protonation states. The optimization of charge-charge interactions via directed mutations has successfully enhanced the thermal stability of different proteins and could be a key to protein engineering improvement. The server presents the electrostatic free energy contribution of each polar-charged residue to protein native state stability. The server also indicates which residues contribute to destabilizing the protein native state with positive energy and the side chain exposed to solvent. This residue is a candidate for mutation to increase protein thermostability as a function of the chosen pH condition. The web server is freely available at UNESP (São Paulo State University - DF/IBILCE): http://tksamc.df.ibilce.unesp.br.

## 1 Introduction

The thermal stability of proteins involves a balance of several interactions. A knowledge of the principals that govern these interactions is fundamental to develop biotechnological applications rationally.^1–3^ Among these interactions, electrostatic effects are widely known to be crucial for thermal stability.^4–9^ Solution conditions, such as salt concentration and pH tend to have a great influence on protein stability.^10–13^

Makhatadze and co-workers have explored the optimization of charge-charge interactions via directed mutations and have successfully enhanced the thermal stability of different proteins.^14–16^ In the present paper, a Web Server that uses a similar method for predicting mutations for the enhancement of thermal stability is presented. This computational method consists in calculating the electrostatic free energy contribution of each ionizable residue of a protein using the Tanford-Kirkwood model with a correction that takes into account the solvent accessibility of these residues (TKSA).^17–19^ All calculations are made for the protein in its native state (see further details in *Material Methods*). The difference between the approach presented here and that used by Makhatadze and co-workers lies in the way the ionizable states of the residues are sampled. In the present study, the charges are sampled by the Monte Carlo method, which was found to be faster than the charge sample made using the Genetic Algorithm.

The TKSAMC is a web server which calculates the protein charge-charge interaction rapidly and consistently. The server provides the residues that are not electrostatically favorable to protein stability (possible target for mutations). The user can also select the temperature and the pH range for the calculations. The web server is freely available at UNESP (São Paulo State University - DF/IBILCE): http://tksamc.df.ibilce.unesp.br.

## 2 MATERIALS AND METHODS

### 2.1 Input files

The main input of this server is a file that contains the coordinates of the atoms of the protein’s native structure. This file can be uploaded in the standard format of Protein Data Bank atom position. A sample of the PDB file can also be downloaded as an example in the server initial page. The most important parameter in TKSA-MC calculation is the pH of a solution; users must choose between a single pH value or a range of pH. If the users opt for a range of pH, the TKSA-MC is calculated for every 0.5 pH unit within the chosen pH range. Another input parameter is temperature, which can be specified or not; the default temperature in this server is 300K. Finally, users can opt to provide an email address to be notified when a given calculation is concluded.

### 2.2 Server calculation - TKSA-MC

In the Tanford-Kirkwood model (TK), the protein is treated as a spherical cavity with dielectric constant, *ϵ_p_* and radius *b*, surrounded by an electrolyte solution modeled according to the Debye-Hückel theory.^17^ Shire *et al* modified this model incorporating a solvent static accessibility correction for each ionizable residue (TKSA model).^18^ This modification was mostly introduced to take into account the irregular protein-solvent interface. The present model is now referred to as the Tanford-Kirkwood model with a solvent accessibility (TKSA). ^18,20,21^ The interaction energy between two charged residues is given by,

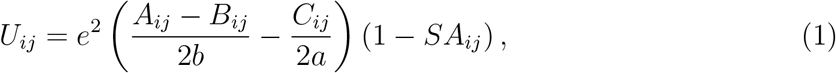

where *e* is the elementary charge; *b* is the radius of the protein; *a* is the closest possible approach distance of an ion; *A_ij_*, *B_ij_* and *C_ij_* are parameters obtained from the analytical solution of the Tanford and Kirkwood model, which are functions of the distance between ionizable residues, the dielectric constants, and the ionic strength;^17,22,23^ and *SA_ij_* is the average of the solvent-accessible surface area of residues *i* and *j*.^18,20,24^

Once the electrostatic energy between the titratable residues has been determined, it is possible to calculate the free energy, Δ*G_N_*(*χ*), for the protein native state in a given state of protonation *χ*:^25,26^

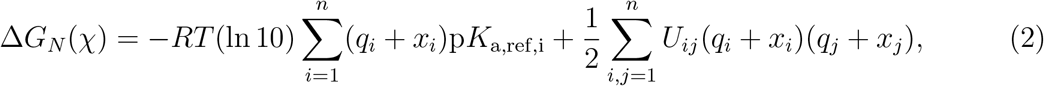

where *R* is the ideal gas constant; *T* is the temperature; *q_i_* is the charge of the ionizable residue *i* in its deprotonated state; *x_i_* is 0 or 1, according to the protonation state of the residue i; and p*K*_a,ref,i_ is the reference *pK_a_*. The probability that the protein will be in its native state with a particular state of protonation, *ρ_N_*(*χ*), is given by,

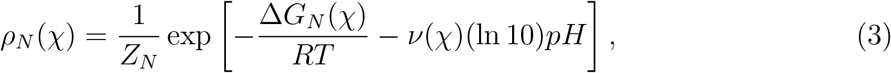

where *ν*(*χ*) is the number of ionizable residues that are protonated in the state of protonation *χ*, and *Z_N_* is the partition function for the protein in its native state. In the method, it is assumed that the unfolded state does not contribute to the electrostatic interactions.^27^ The protonation states *χ* are calculated via Metropolis Monte Carlo. The mean total electrostatic energy, 〈*W_qq_*〉 is computed by the average over these protonation states, taking into account Eq. 3:

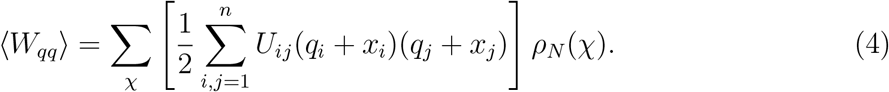

Ibarra-Molero and Makhatadze have shown that the contribution of electrostatic interaction energy to free energy, Δ*G_qq_*, can be described by Δ*G_qq_* ≈ −〈*W_qq_*〉.^26^

### 2.3 Output files

For each submitted job a report is generated. The result page has a specific link which is sent to the e-mail provided by the user. All the data used in the method calculation and the plotted figures can be downloaded as a compressed folder in a link at the top of the page. An output file with the description of the charge positions, the reference *pK_a_*, the normalized solvent-accessibility surface area, the energy contribution of each residue and the total energy is also available. If the submitted job is a single pH run, the result page presents a Δ*G_qq_* bar graph, like that shown in Figure 1. If the submitted job uses the pH range calculation option, the result page gives the Δ*G_qq_* bar graph at the top and the Δ*G_elec_* as a function of pH at the bottom, as shown in Figure 2.

**Figure 1:**
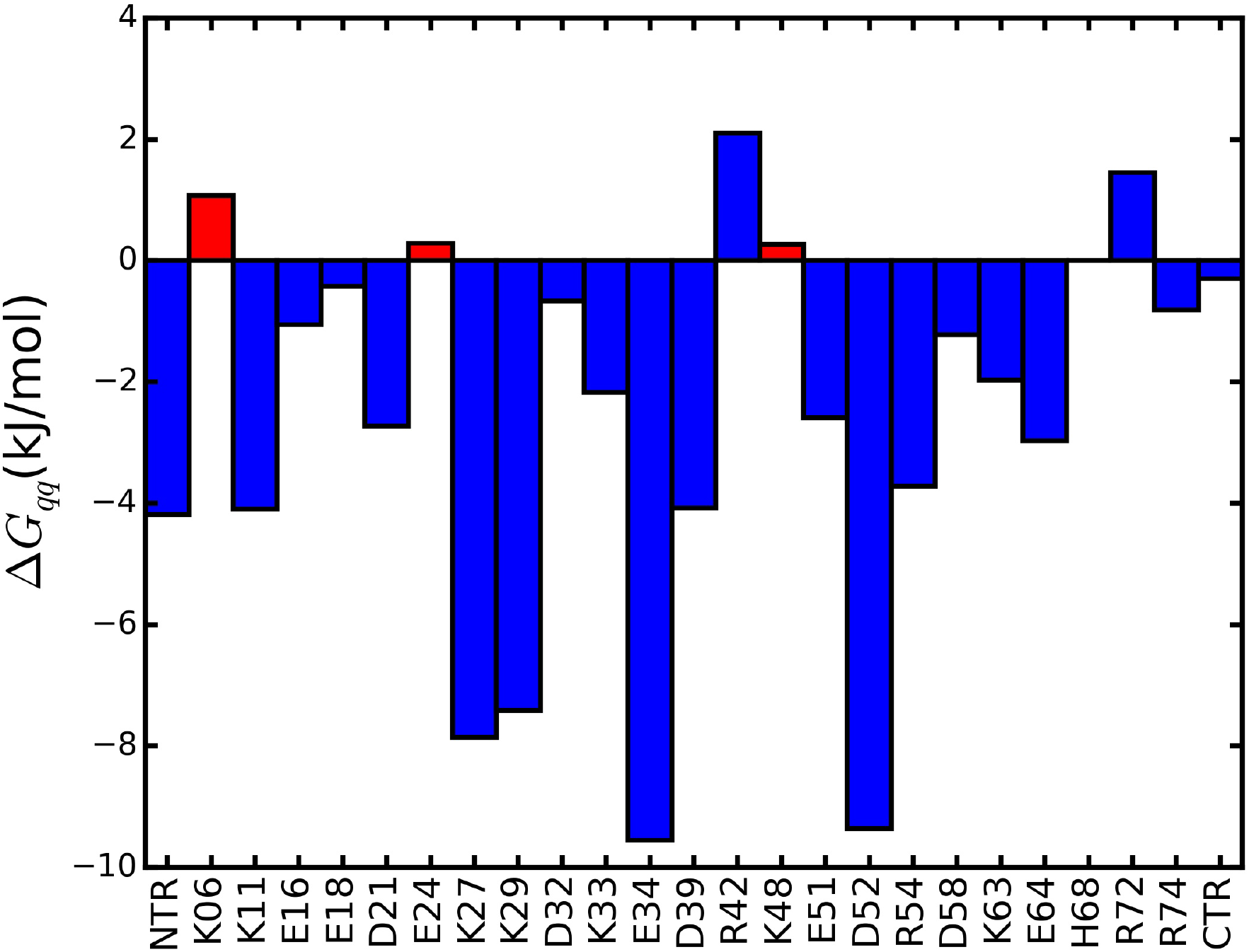
Charge–charge interaction energy Δ*G_qq_* calculated by the TKSA-MC model for each ionizable residue, in this example, for ubiquitin. The simulation parameter values were pH 5.5 and temperature 300K. The ubiquitin structure used was PDB ID: 1UBQ. It has 25 ionizable groups, in which NTR and CTR are the N and C terminals, respectively.

**Figure 2:**
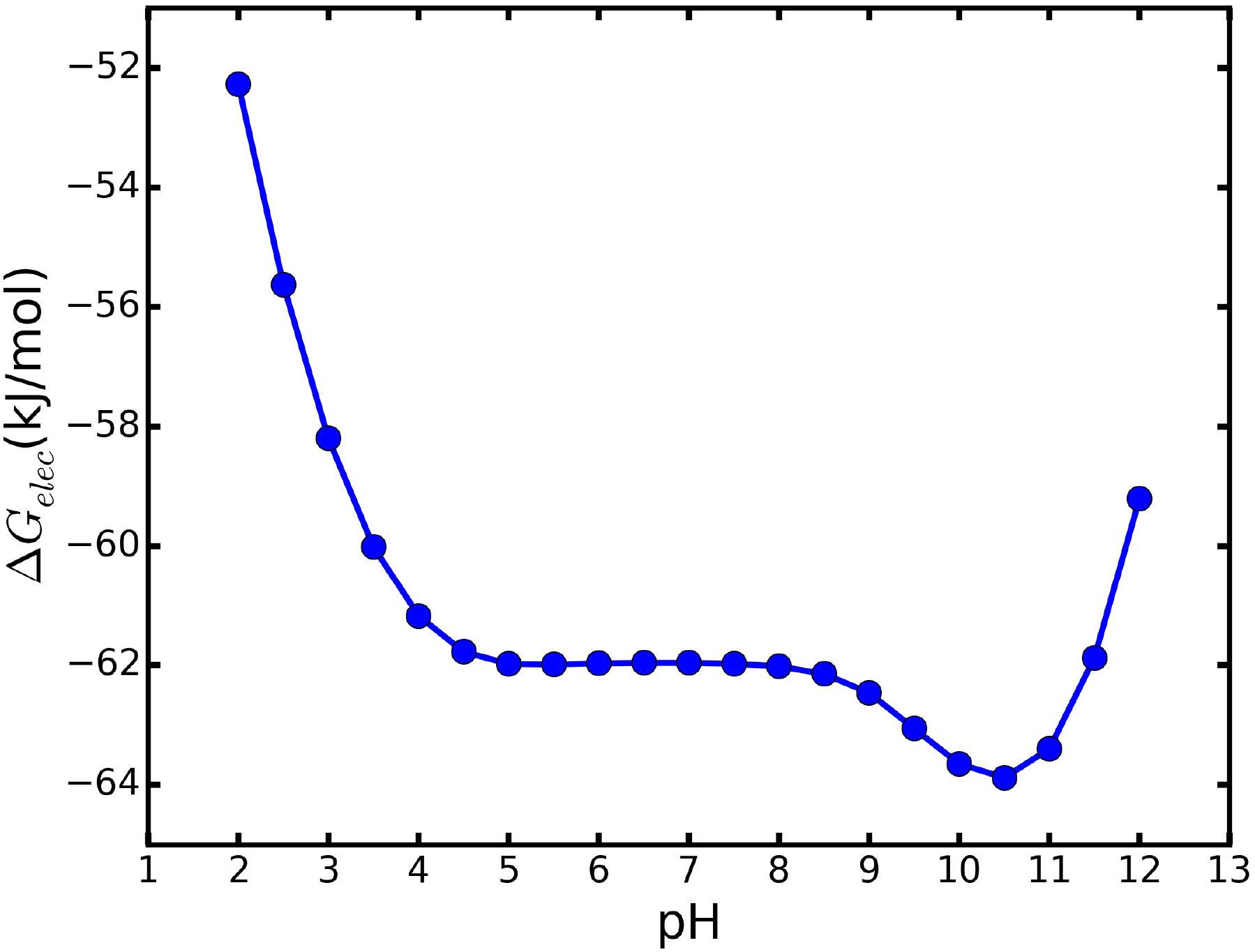
Δ*G_elec_* as a function of pH calculated from pH 2 to 12 for ubiquitin in a total of 21 TKSA-MC runs at a temperature of 300K.

## 3 RESULTS

### 3.1 Δ*G_qq_* bar graphs analysis

The TKSA-MC method indicates the residues which contribute to destabilizing the protein native state. The algorithm calculates the protein electrostatic energy, taking into account the contribution of each residue with a polar-charged side chain. Figure 1 presents an example of the Δ*G_qq_* bar graph for ubiquitin.^27^

The bars indicate the charge-charge energy contribution of each ionizable residue to protein native state stability. The red bars indicate the candidate residues to be mutated to increase protein thermostability. The selection criteria follow the methodology introduced by Ibarra-Molero et al.,^26^ which indicates the residues that present unfavorable energy values Δ*G_qq_* ≥ 0 and are exposed to solvent with SASA ≥ 50%. In the ubiquitin example, the highlighted residues are K06, E24 and K48, and this information is also printed on the result screen page with its respective Δ*G_qq_* and SASA values. All the data for plotting the Δ*G_qq_* bar graph figure is available for download in the output files. The total energy contribution to protein native state stability is given as a sum of the Δ*G_qq_* for each ionizable group, named as Δ*G_elec_*. The Δ*G_elec_* the ubiquitin in pH 5.5 is −61.99kJ/mol.

### 3.2 Δ*G_elec_* and pH dependence

The total energy contribution Δ*G_elec_* as well as the ionization degree of each polar side chain residue and its Δ*G_qq_* are pH dependent. Figure 2 presents an example of Δ*G_elec_* as a function of pH for ubiquitin. The pH range in the example shown in Figure 2 was from 2 to 12 in steps of 0.5 in a total of 21 TKSA-MC runs. All the data used to plot the Δ*G_elec_* curve and the respective pH bar graph figures are available for download in the output files.

## 4 CONCLUDING REMARKS

The TKSA-MC method is a Monte Carlo/Metropolis based approach to sampling the ionizable protein groups. This method is an adaptation of the genetic algorithm sampling approach TKSA-GA method introduce by Ibarra-Molero et al..^26^ The web server provides a fast calculation of the charge-charge interaction using the Tanford-Kirkwood model protein approach with pre-selected pH and temperature values.

As a result, the TKSA-MC indicates the residues which contribute to destabilizing the protein native state. The charge-charge interaction optimization by mutation is applied to protein engineering.^9,16,26–29^ The indicated residue is a candidate for mutation for increasing the protein thermostability in the chosen pH and temperature conditions. It is recommended that the TKSA-MC also be run for the mutated protein, and the pH range option be run to check the pH-dependence of the protein stability due to electrostatic contribution Δ*G_elec_*. The server also has a section with a discussion of the expected results and how to interpret them, as well a section containing the theory and methodology details. A personalized support via e-mail is also provided.

